# Profiling of the β-glucosidases identified in the genome of *Penicillium funiculosum*: Insights from genomics, transcriptomics, proteomics and homology modelling studies

**DOI:** 10.1101/2022.10.18.512808

**Authors:** Omoaruemike Ebele Okereke, Mayank Gupta, Olusola A. Ogunyewo, Kanika Sharma, Syed Shams Yazdani

## Abstract

Enzymatic lignocellulosic biomass conversion to bioethanol is dependent on efficient enzyme systems with β-glucosidase as a key component. In this study, we performed in-depth profiling of the various β-glucosidases present in the genome of the hypercellulolytic fungus; *Penicillium funiculosum* using genomics, transcriptomics, proteomics and molecular dynamics simulation approaches. Of the eight β-glucosidase genes identified in the *P*. *funiculosum* genome, three were found to be extracellular, as evidenced by presence of signal peptides and mass spectrometry. Among the three secreted β-glucosidase, two belonged to the GH3 and one belonged to GH1 families. Modelled structures of these proteins predicted a deep and narrow active site for the GH3 β-glucosidases (*Pf*Bgl3A and *Pf*Bgl3B) and a shallow open active site for the GH1 β-glucosidase (*Pf*Bgl1A). The enzymatic assays indicated that *P. funiculosum* secretome showed high β-glucosidase activities with prominent bands on 4-methylumbelliferyl β-D-glucopyranoside (MUG) zymogram. To understand the contributory effect of each of the three secreted β-glucosidases (*Pf*Bgls), the corresponding gene was deleted separately and the effect of the deletion on β-glucosidase activity of the secretome was examined. Although not the most abundant β-glucosidase, *Pf*Bgl3A was found to be the most significant one as evidenced by a 42 % reduction in β-glucosidase activity in the Δ*Pf*Bgl3A strain. To improve the thermostability, two mutants of *Pf*Bgl3A were designed with the help of molecular dynamics (MD) simulation and were expressed in *Pichia pastoris* for evaluation. The *Pf*Bgl3A mutant (Mutant A) gave 1.4 fold increase in the half-life (T_1/2_) of the enzyme at 50°C.

**IMPORTANCE:** Commercially available cellulases are majorly produced from *Trichoderma reesei*. However, external supplementation of the cellulase cocktail from this host with exogenous β-glucosidase is often required to achieve desired optimal saccharification of cellulosic feedstocks. This challenge has led to exploration of other cellulase-producing strains because of the importance of this class of enzymes in the cellulose deconstruction machinery. The non-model hypercellulolytic fungus *Penicillium funiculosum* has been studied in recent times and identified as a promising source of industrial cellulases. Various genetic interventions targeted at strain improvement for cellulase production have been performed. However, the β-glucosidases of this strain have remained largely understudied. This study, therefore, reports profiling of all the eight β-glucosidases of *P. funiculosum* via molecular and computational approaches and enhancing thermostability of the most promising β-glucosidase via protein engineering. The results of this study set the background for future engineering strategies to transform the fungus into an industrial workhorse.

## INTRODUCTION

Industrial processes are being harmonized to be more environment-friendly systems. In the energy sector, researchers are searching for alternative cleaner sources of energy to reduce pollution caused by fossil fuels. This has resulted in increased research into the production of second-generation biofuels from lignocellulosic materials, which represent an abundant, viable, cheap, and clean alternative (1–5). Lignocellulosic biomass is considered to be the most abundant renewable bioresource on earth. Their abundance and renewability make them an ideal raw material for the production of second-generation biofuels (6). The conversion process of lignocellulosic biomass to bioethanol includes several steps, such as pretreatment, enzymatic hydrolysis, fermentation and product recovery. These steps require a combination of an efficient, cost-effective and sustainable strategy in order to achieve high bioethanol yields at a reduced cost (7, 8). Currently, the biofuel production cost is much higher as compared to fossil fuel cost and, thus, there is a persistent effort in research and development to ensure market level availability of lignocellulosic biofuel (9). The bioconversion of lignocellulosic biomass to fermentable sugars is facilitated by cellulolytic enzymes. The high cost of these enzymes has been one of the major limitations to the commercialization of lignocellulosic ethanol (10).

The cellulase enzymes acting on the cellulose portion of lignocellulosic biomass majorly consist of three types of enzymes: endo-β-1, 4-glucanase (EG), exo-β-1, 4-glucanase or cellobiohydrolase (CBH) and β-glucosidase (BGL). Complete enzymatic hydrolysis of cellulose requires a synergistic action of these three cellulases (11). EG acts on the cellulose chain randomly yielding cello-oligosaccharides, while CBH cleaves the reducing and non-reducing ends of the chain releasing cellobiose and cello-oligosaccharides. These products are the substrates for BGL which then cleaves them to glucose. Accumulation of cellobiose and cello-oligosaccharides leads to inhibition in the activities of EG and CBH. BGL thus relieves this inhibition allowing increased glucose production (12). Cellulases are inducible enzymes synthesized by a number of microorganisms when growing on cellulosic materials (13). The majority of these organisms are bacteria, however, fungi are the preferred source because of the ease of recovery, higher thermostability as well as higher activity of these enzymes. Currently, commercial cellulases are usually produced from *Trichoderma reesei*, although this is supplemented with β-glucosidase from *Aspergillus spp*. because *Trichoderma reesei* cellulase cocktail is deficient in β-glucosidase (12). In recent times, some other fungi from the *Penicillium* and *Aspergillus* genera have been used to generate lignocellulolytic enzymes (10). *Penicillium* strains in particular have been presented as a potent alternative to the *T. reesei* cellulase cocktail and there has been increasing documentation of the production of industrially viable cellulase enzyme cocktails from *Penicillium* strains which have been shown to be richer in β-glucosidase than *Trichoderma reesei* (14, 15). Recently, Ogunmolu *et al* (16) identified a strain of the filamentous fungus *Penicillium funiculosum* (NCIM1228) which has outstanding biomass hydrolyzing potential comparable to *Trichoderma reesei*. The organism was found to be a hypercellulolytic fungus and in order to develop NCIM1228 into an industrial workhorse, several genetic interventions have been carried out to deregulate and overexpress the cellulolytic genes to improve cellulase production, activity and stability of the enzymes, resulting in increased catalytic activity, expression and secretion of both exo- and endo-cellulases (17–21). However, little has been studied for the enhancement of the activity and thermostability of *P. funiculosum* β-glucosidase. Since β-glucosidase is the rate-limiting enzyme in the cellulase mediated hydrolysis of lignocellulolytic biomass catalyzing the final step, it plays a key role in the determination of the hydrolytic efficiency of the enzyme cocktail. Additionally, β-glucosidases are usually less stable than exo-glucanases and endo-glucanases and are particularly prone to inactivation caused by temperature changes during hydrolysis. This is a major setback that greatly impacts negatively the efficiency of the enzymatic hydrolysis of lignocellulose. Therefore, there is a need for genetic interventions to improve the efficiency of the β-glucosidase enzyme which will invariably improve the *P. funiculosum* lignocellulosic biomass hydrolytic activity. However, for successful genetic interventions, a knowledge of the enzyme profile, expression levels, individual contribution to the overall enzyme activity, and thermostability is essential for strategic design. This study was therefore aimed to determine the β-glucosidase profile of a catabolite de-repressed *Penicillium funiculosum* NCIM1228 (PfMig1^88^) with a view to evaluating the abundance and contribution of individual β-glucosidases to the overall saccharification efficiency of the fungal secretome. The most efficient β-glucosidase was then modeled to engineer the thermostability features and expressed in *Pichia pastoris* to evaluate the kinetic parameters and thermostability.

## RESULTS AND DISCUSSION

### Sequence and phylogenetic analysis of *Penicillium funiculosum* β-glucosidases

Multiple genes are known to encode β-glucosidases in cellulolytic organisms. Genome sequencing and annotation of *Penicillium funiculosum* NCIM1228 in our laboratory revealed the presence of eight β-glucosidase genes coding for β-glucosidases with molecular weights varying from 54.6 to 90.8 kD and amino acid lengths of 480 - 855 aa (Table 2). This is consistent with other documented reports on multiple β-glucosidases identified in various organisms (22–25). The sequences of these *Penicillium funiculosum* β-glucosidases (*Pf*Bgls) (Supplementary Fig. S1) were deposited in GenBank with accession numbers OM595351 to OM595358. Sequence analysis of the eight *Pf*Bgls using Signal P software predicted the presence of signal peptide in the ORF of *Pf*Bgl3A, *Pf*Bgl3B and *Pf*Bgl1A at amino acid positions 19 & 20, 18 &19 and 16 &17, respectively (Supplementary Fig. S2) and are therefore designated as the three extracellular *Pf*Bgls in *P. funiculosum*. No signal peptides were predicted for the other five *Pf*Bgls. Domain prediction (Supplementary Fig. S3) revealed that the *Pf*Bgls had conserved domains documented for GH3 and GH1 β-glucosidases. The glycoside hydrolase family 3 superfamily domain (pfam01915 and pfam00933) were present in *Pf*Bgl3A and *Pf*Bgl3B confirming that they belong to the GH3 CAZy family. They also had a fibronectin type III-like domain (pfam14310) and a Bgl1X domain associated with carbohydrate transport and metabolism. *Pf*Bgl1A on the other hand is a much smaller protein with one glycosyl hydrolase family 1 superfamily catalytic domain (pfam00232) placing it in the GH1 CAZy family. Similar domains have been reported for GH3 and GH1 proteins (24, 26). Since *Pf*Bgl3A, *Pf*Bgl3B and *Pf*Bgl1A were the only ones predicted to have a secretary signal sequence and thus most likely to be secreted extracellularly to contribute to biomass hydrolysis, they were selected for further analysis. NCBI BLAST of these three extracellular β-glucosidases of *P. funiculosum* showed greater than 90% similarity with some well-documented β-glucosidases of the *Penicillium* family. Phylogenetic analysis of *Pf*Bgl3A, *Pf*Bgl3B and *Pf*Bgl1A puts them into three major clades, with *Pf*Bgl3A and *Pf*Bgl3B evolutionally related to GH3 family proteins while the third clade has *Pf*Bgl1A and GH1 β-glucosidases (Fig. 1). All three proteins showed the highest similarity with β-glucosidases from *Penicillium occitanis* and *Talaromyces* spp.

**Fig. 1.**
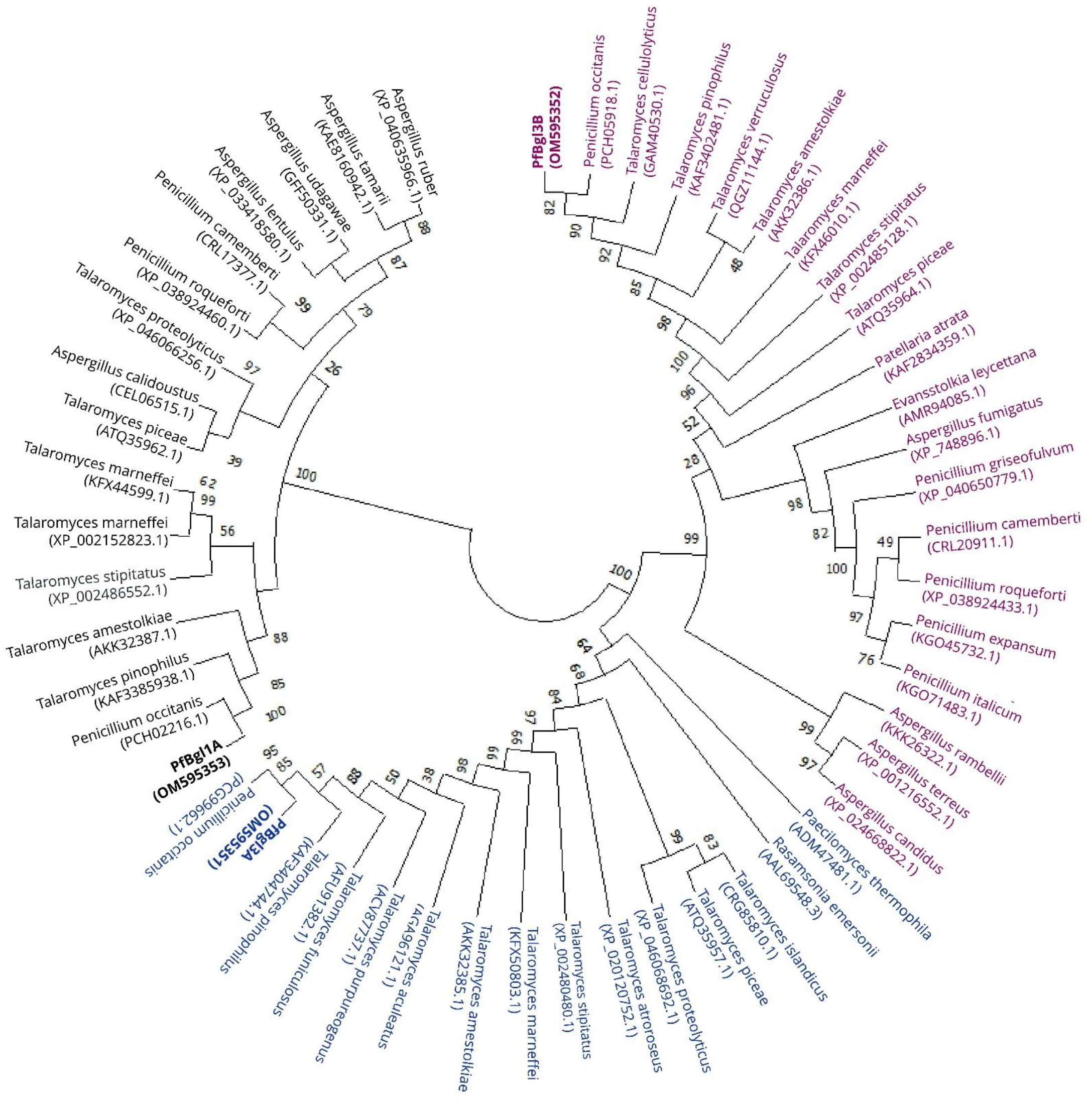
Phylogenetic analysis of *Penicillium funiculosum* NCIM1228 β-glucosidases (*Pf*Bgl3A, *Pf*Bgl3B and *Pf*Bgl1A) by the neighbor-joining method. Sequences used to construct the tree were retrieved from NCBI database. Different colours represent each major clade and numbers represent the bootstrap replicate percentage (1000 replicates)

**Table 1.**
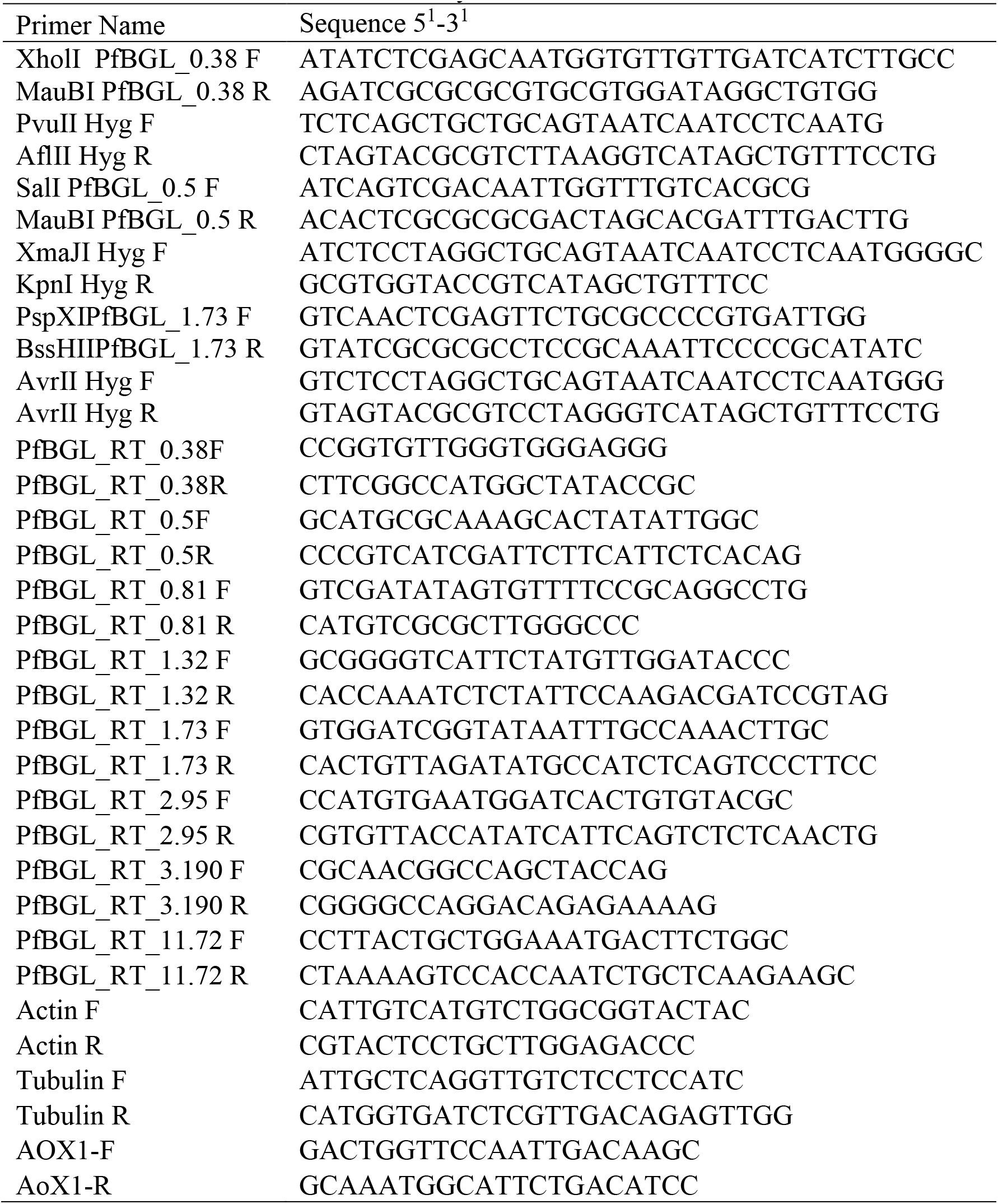
List of Primers used in this study.

**Table 2.**
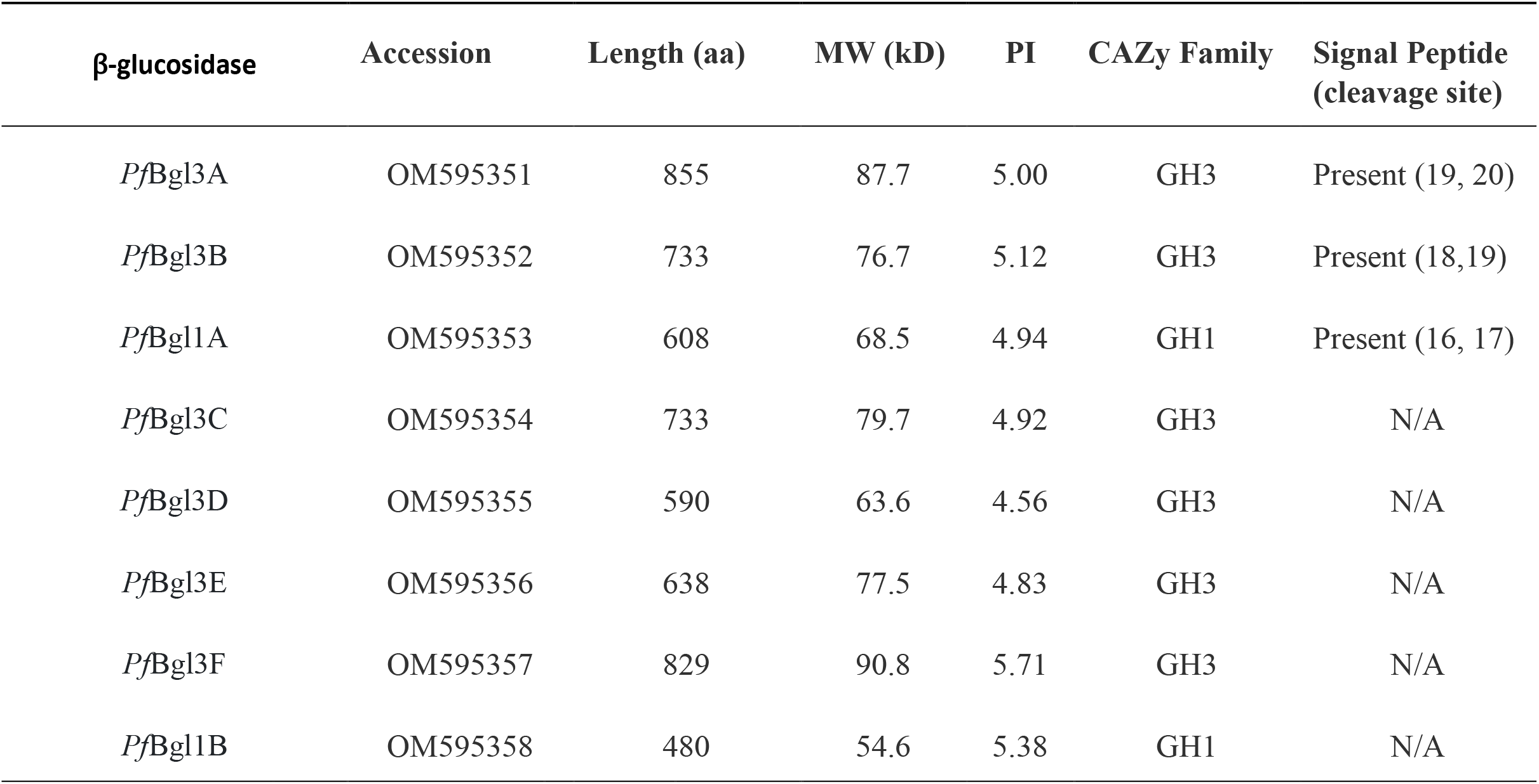
β-glucosidases identified in the genome of *Penicillium funiculosum*.

| β-glucosidase | Accession | Length (aa) | MW (kD) | PI | CAZy Family | Signal Peptide (cleavage site) |
| --- | --- | --- | --- | --- | --- | --- |
| *PfBgl3A* | OM595351 | 855 | 87.7 | 5.00 | GH3 | Present (19, 20) |
| *PfBgl3B* | OM595352 | 733 | 76.7 | 5.12 | GH3 | Present (18,19) |
| *PfBgl1A* | OM595353 | 608 | 68.5 | 4.94 | GH1 | Present (16, 17) |
| *PfBgl3C* | OM595354 | 733 | 79.7 | 4.92 | GH3 | N/A |
| *PfBgl3D* | OM595355 | 590 | 63.6 | 4.56 | GH3 | N/A |
| *PfBgl3E* | OM595356 | 638 | 77.5 | 4.83 | GH3 | N/A |
| *PfBgl3F* | OM595357 | 829 | 90.8 | 5.71 | GH3 | N/A |
| *PfBgl1B* | OM595358 | 480 | 54.6 | 5.38 | GH1 | N/A |

### Structural analysis and computational evaluation of substrate interactions of *Pf*Bgl3A, *Pf*Bgl3B and *Pf*Bgl1A

Structural models of the three extracellular β-glucosidases of *P. funiculosum* generated by homology modelling revealed distinct proteins with *Pf*Bgl3A and *Pf*Bgl3B having the characteristic (α/β)_8_ triose phosphate isomerase (TIM) barrel and (α/β)6 sandwich domains connected by linkers, while *Pf*Bgl1A has one domain with the (β/α)_8_ barrel type structure (Fig. 2). The modeled structures were validated with Ramachandran plots (27) using PROCHECK module of PDBsum server. Ramachandran analysis indicated that 99.6 %, 99.5 % and 97.5 % of the amino acid residues of *Pf*Bgl3A, *Pf*Bgl3B and *Pf*Bgl1A respectively are in the favoured and allowed regions (Fig. 2). A comparison of the surface structures (Fig 3) showed that the active sites of *Pf*Bgl3A and *Pf*Bgl3B are open and shallow, whereas that of *Pf*Bgl1A is deep, narrow and tunnel-like consistent with reports on active site configuration for GH1 β-glucosidases (26, 28–31). This active site conformation is thought to improve the glucose tolerance of GH1 proteins by preventing the direct access of glucose to the active site (32). To determine the amino acid residues implicated in substrate binding at the active site, the *Pf*Bgls were docked with cellobiose. *Pf*Bgl3A, *Pf*Bgl3B and *Pf*Bgl1A formed six, eight and seven hydrogen bonds respectively with cellobiose (Fig 3), with binding energies ranging from −6.3 to −6.9 kcal/mol. Binding energies in the range of −4.9 to −8 kcal/mol have been reported for β-glucosidases from *Aspergillus fumigatus* and *Paenibacillus polymyxa* (24, 33). Yadav et al. (34) however reported a much lower binding energy of −11.33 to −13.29 kcal/mol for *Paenibacillus lautus*. The amino acid residues forming hydrogen bonds in the binding pocket of *Pf*Bgl3A are Asp 273, Arg 196, Arg 152, Glu 502 and Arg 94, while hydrogen bonds are formed by Ser 405, Lys 172, Asp 250, Asp 75, Glu 462, Arg 81 and Tyr 82 in that of *Pf*Bgl3B. Previously, Ramani et al., (35) also reported the catalytic acid/base residue Glu 502 and nucleophile Asp 273 in the catalytic site of a β-glucosidase from *Penicillium funiculosum* NCL1 strain consistent with our results for *Pf*Bgl3A. Amino acid residues forming hydrogen bonds in the catalytic pocket of *Pf*Bgl1A are Glu 567, Gln 165, Tyr 325, Asp 432 and Glu 321. Here the catalytic acid/base and nucleophile residues are Glu as reported for GH1 family β-glucosidases.

**Fig. 2.**
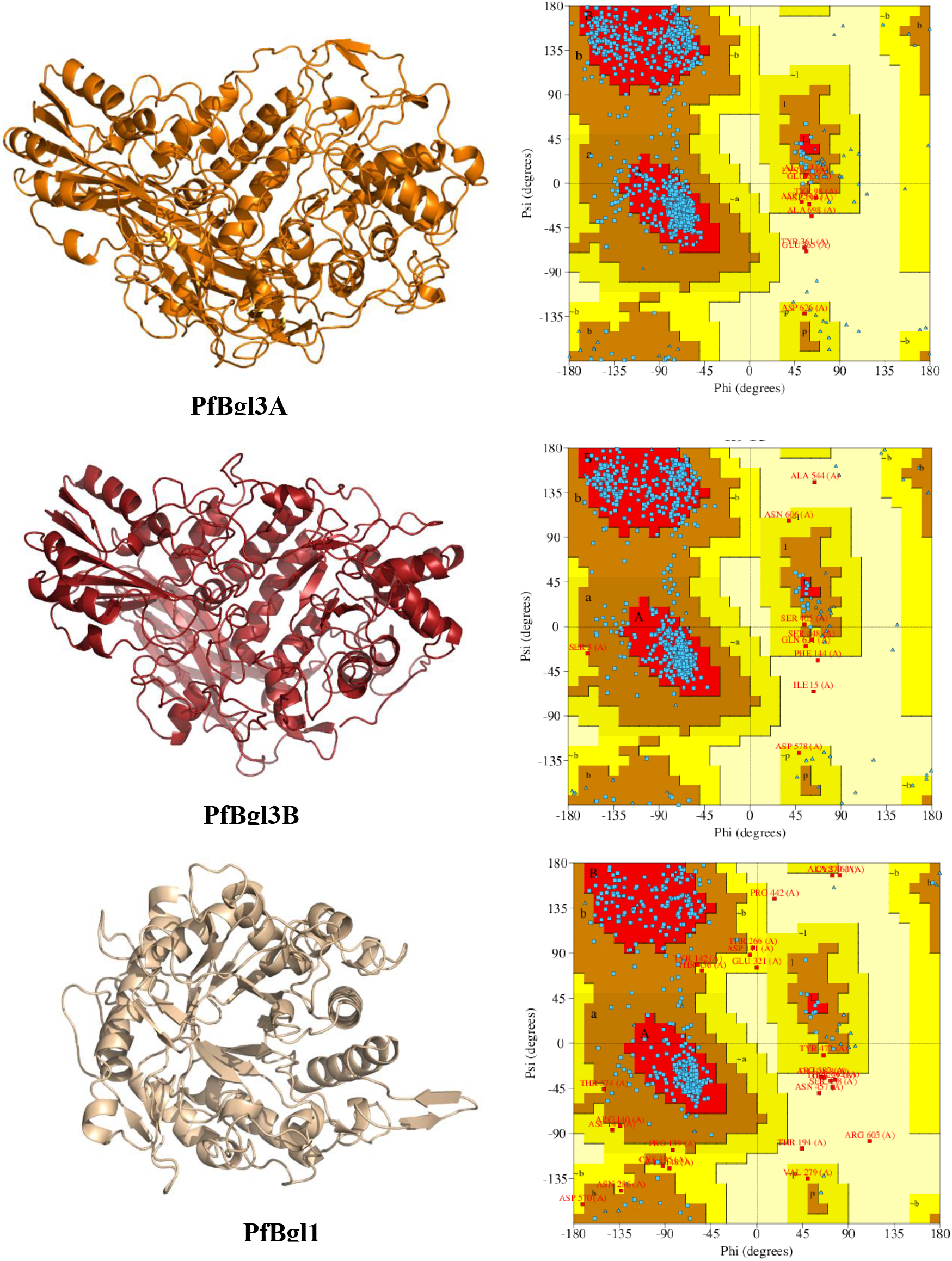
Ribbon diagram representation (left panel) and Ramachandran plots (right panel) of the predicted structures of the extracellular *Pf*Bgls. In the Ramachandran plots, most favoured regions (A, B, L) are coloured red, additionally allowed regions (a, b, l, p) are brown, generously allowed regions (~a, ~b, ~l, ~p) are cream coloured.

**Fig. 3.**
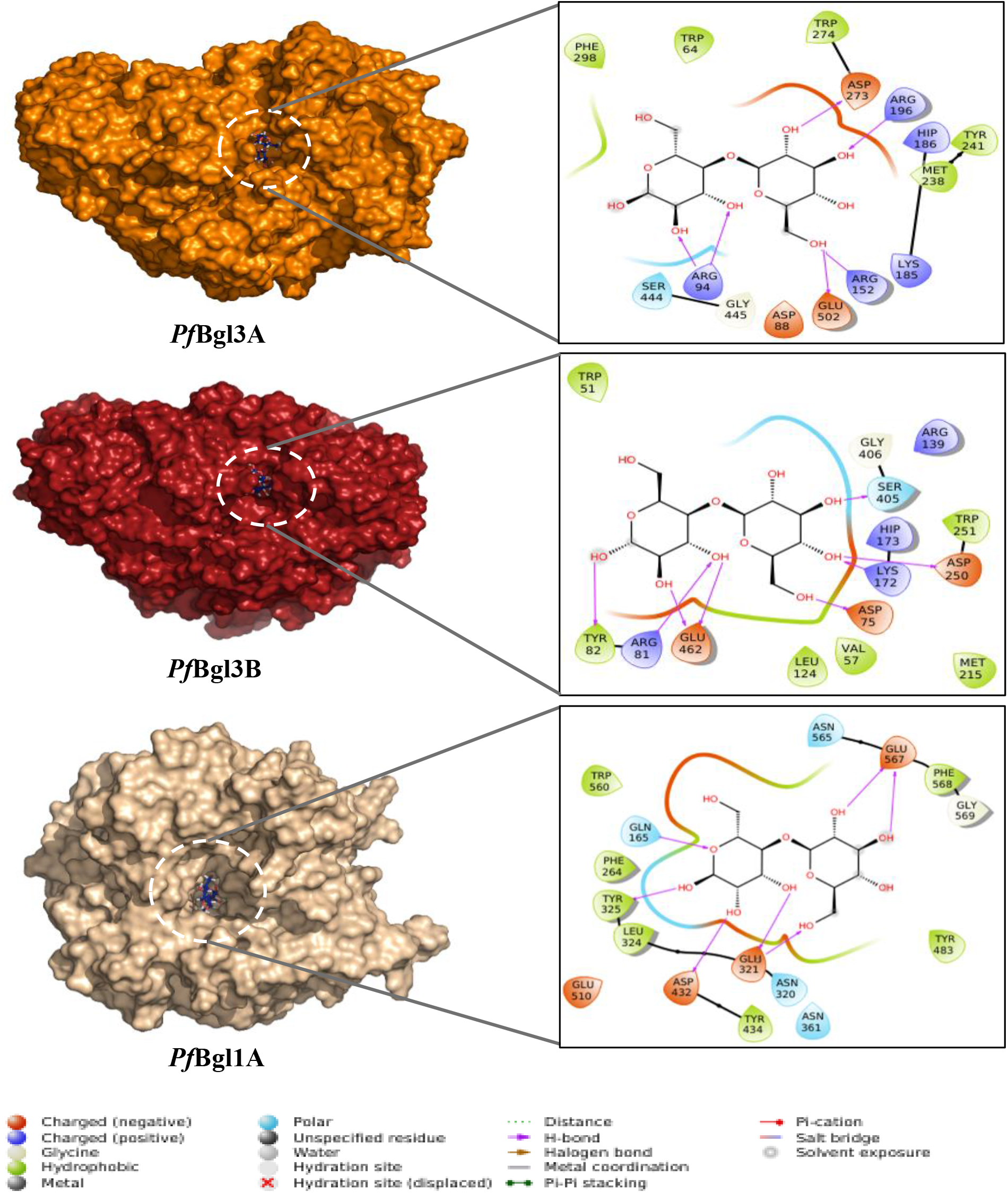
Comparison of the surface structures of *Pf*Bgl models complexed with cellobiose (highlighted with dotted circles).Surrounding amino acid residues of the active sites and ligand interactions are represented in the expanded boxes.

### Evaluation of β-glucosidase activity via MUG Zymography

The presence of active β-glucosidases in *Pf*Mig1^88^ was confirmed by MUG Zymography. The Zymogram (Fig. 4) revealed two prominent distinct bands with intensity reducing with time corresponding to activity loss (Supplementary Table S1). The intensity of the bands indicates that the β-glucosidases of *P. funiculosum* are highly active. There is a correlation between the band intensity and activity of the protein (36) demonstrated by Li et al., (37). They showed that a recombinant *Trichoderma reesei* RUT C30 with enhanced β-glucosidase activity had more prominent bands on MUG gel than the parent RUT C30 strain. We can therefore conclude that *P. funiculosum* secretes highly active β-glucosidases.

**Fig. 4.**
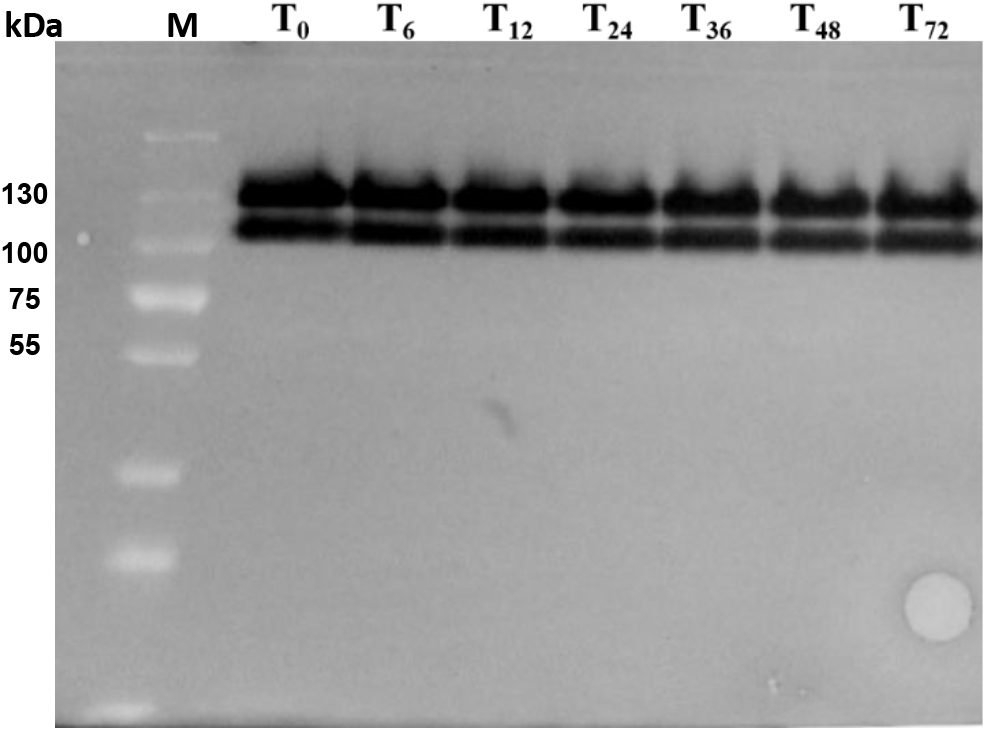
MUG zymograph of PfMig1^88^ secretome. Two distinct bands are visible confirming the presence of highly active β-glucosidases. T0 –T72 are time points corresponding to 0 – 72 h respectively.

### Expression level analysis of *P. funiculosum* β-glucosidases

To determine the expression levels of each of the secreted *P. funiculosum* β-glucosidases, quantitative polymerase chain reaction (qPCR) and proteomics were performed to investigate the transcript abundance of *Pf*Bgl genes and proteins in the secretome, respectively. The quantification cycle values for each gene was normalized with the highly expressed housekeeping gene encoding actin. Data generated revealed that *Pf*Bgl3B had the highest transcript levels when grown in cellulose inducing media containing Avicel and Wheat bran (Fig. 5A). This result correlated with that of the protein expression profile where *Pf*Bgl3B was the most abundant β-glucosidase found in the secretome (Fig 5B, Supplementary Table S2). *Pf*Bgl3A and *Pf*Bgl1A were also significantly abundant in the secretome and although the transcript levels of *Pf*Bgl3A was among the lowest, the protein abundance was relatively high which could be an indication of high translational efficiency. The intracellular β-glucosidases that were found in the secretome were present in comparatively negligible amounts and could be attributed to leakage into the secretome as a result of cell death and lysis. Similar findings were reported by Ogunyewo et al. (38) for *Pf*Mig1^88^ grown in cellulase-inducing media containing pretreated sugarcane bagasse. Our results thus established that the three most abundant and significant β-glucosidases in *P. funiculosum* secretome which contribute to the overall β-glucosidase activity of the fungus are *Pf*Bgl3A, *Pf*Bgl3B and *Pf*Bgl1A.

**Fig. 5.**
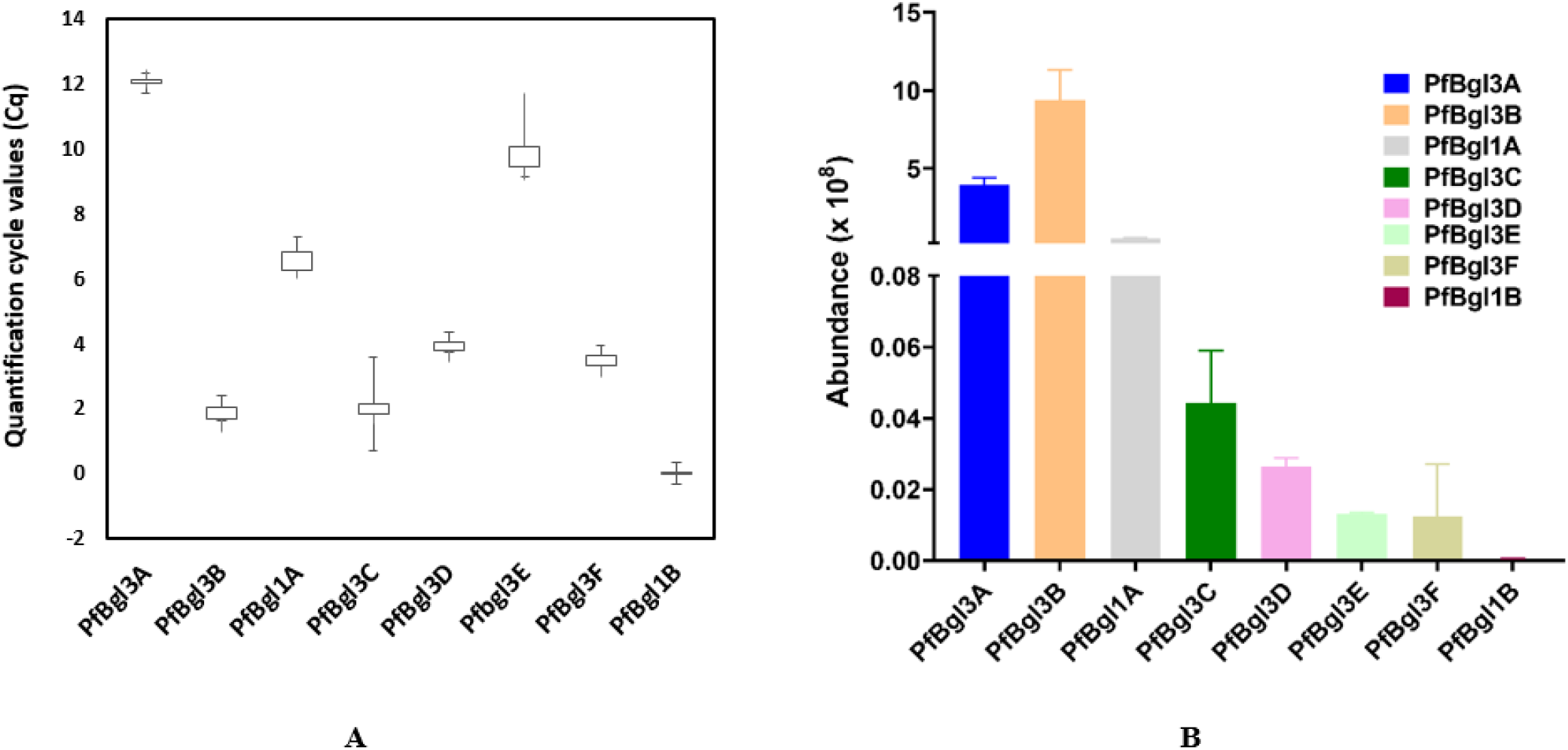
Expression levels of *P. funiculosum* β-glucosidases. (A) Abundance of *Pf*Bgl transcripts. (B) Abundance of β-glucosidases in *P. funiculosum* secretome.

### Functional role of β-glucosidases in *P. funiculosum*

Having established the significant β-glucosidases in *P. funiculosum*, the contribution of each of the *Pf*Bgls to the overall β-glucosidase activity of *P. funiculosum* secretome was investigated. Each of the genes for the extracellular β-glucosidases was deleted creating *Pf*Bgl null strains designated as Δ*Pf*Bgl3A, Δ*Pf*Bgl3B and Δ*Pf*Bgl1A mutants. Our results (Fig. 6A) indicated that deletion of *Pf*Bgl3A and *Pf*Bgl1A resulted in a reduction in β-glucosidase activity by about 42 % and 13% respectively when compared to the native PfMig1^88^. However, the deletion of *Pf*Bgl3B did not lead to any significant change in the β-glucosidase activity of the *P. funiculosum* secretome. This was surprising considering high abundance of both transcript and protein for *Pf*Bgl3B, and thus the deletion mutant may need to be thoroughly analyzed further, although similar observations have been made (39–41). Since this study aims to explore β-glucosidases of PfMig1^88^ with a view to improving the saccharification efficiency of the secretome, we also analyzed the effect of the deletion of each of these genes on the glucose yield from the hydrolysis of pretreated sugarcane bagasse at 15 % solid load and 5 FPU/g biomass protein concentration. This was to determine the contributive effect of each to the overall saccharification efficiency. We observed that deletion of *Pf*Bgl3A resulted in significant reduction (29%) in glucose yield. On the other hand, deletion of *Pf*Bgl3B showed non-significant reduction (i.e., 3.6%), while deletion of *Pf*Bgl1A showed only 6.8% reduction in glucose yield from sugarcane bagasse saccharification (Fig. 6B). Our findings have therefore highlighted the roles of *P. funiculosum* β-glucosidases in biomass hydrolysis. It is evident from our findings that *Pf*Bgl3A is a significant β-glucosidase in *P. funiculosum* as its deletion led to 42% decreased in overall β-glucosidase activity and 29% decrease in glucose release from biomass and could be a viable target for strain improvement aimed at enhancing saccharification efficiency.

**Fig. 6.**
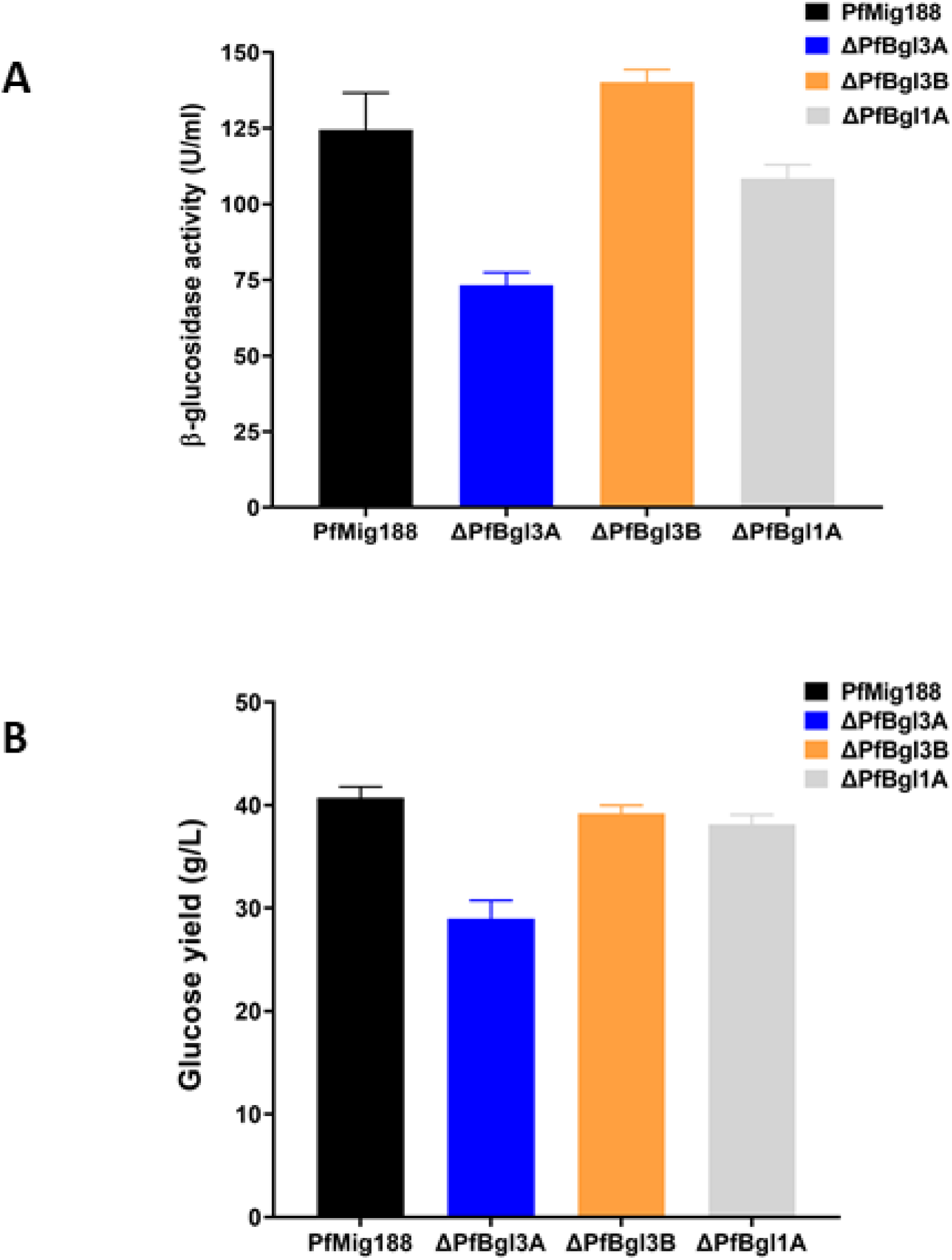
Comparison of the enzyme activities of *Pf*Bgl deletion mutants. (A) β-glucosidase activity measured by release of p-nitrophenol from p-nitrophenol-β-D-glucopyranoside. (B) Glucose yield from saccharification of sugarcane bagasse at 15 % solid loading and protein concentration of 5FPU/g biomass.

### Engineering and expression of thermostable variants of *Pf*Bgl3A

After establishing *Pf*Bgl3A as a potential target for genetic modifications, we chose to design variants of the protein with a view to improving the overall thermostability of β-glucosidase in the enzyme concoction from *P. funiculosum*. Many reports usually target improving the thermostability at elevated temperatures (42) however, our focus was to improve the stability at 50 °C which is the optimum temperature for biomass hydrolysis. We employed the rational approach design strategy; an established method that has been used to design thermostable and glucose-tolerant β-glucosidase mutants (43–45). Initial MD simulation on *Pf*Bgl3A gave an indication of the stability of the protein and amino acid residues with highest fluctuations (Fig. 7A). From root mean square deviation (RMSD) and root mean square fluctuation (RMSF) values, points of highest fluctuations were selected for mutations based on data from multiple sequence analysis with sequences of thermostable β-glucosidases having structures reported in PDB (Fig. 8). MD simulation results after incorporating the proposed amino acid substitutions (Fig. 7B) gave an indication of mutations which lowered the RMSF and RMSD values. These were chosen and the two mutants (mutant A and mutant B) designed and expressed in *P. pastoris*. SDS –PAGE (Fig. 9A) confirmed that the proteins were indeed expressed however, the molecular weights were higher than the theoretical amino acid sequence-deduced molecular weight which could be attributed to changes in glycosylation pattern by *P. pastoris* (46). Furthermore, our results indicated that mutant A had improved thermostability (Fig. 9B) giving 1.4 fold increase in half-life (T1/2) as compared to *Pf*Bgl3A (Table 3). Amino acid substitutions was done in loop regions away from the active site in order not to interfere with the hydrolytic efficiency of the protein and our results showed that hydrolytic activity of mutant A is similar to that of the native strain. Mutant B however had significantly reduced thermostability and hydrolytic activity (Fig 9 B and C). This could be attributed to the number of amino acid substitutions being too many which may have interfered with the secondary structure and consequently the function of the protein.

**Fig. 7.**
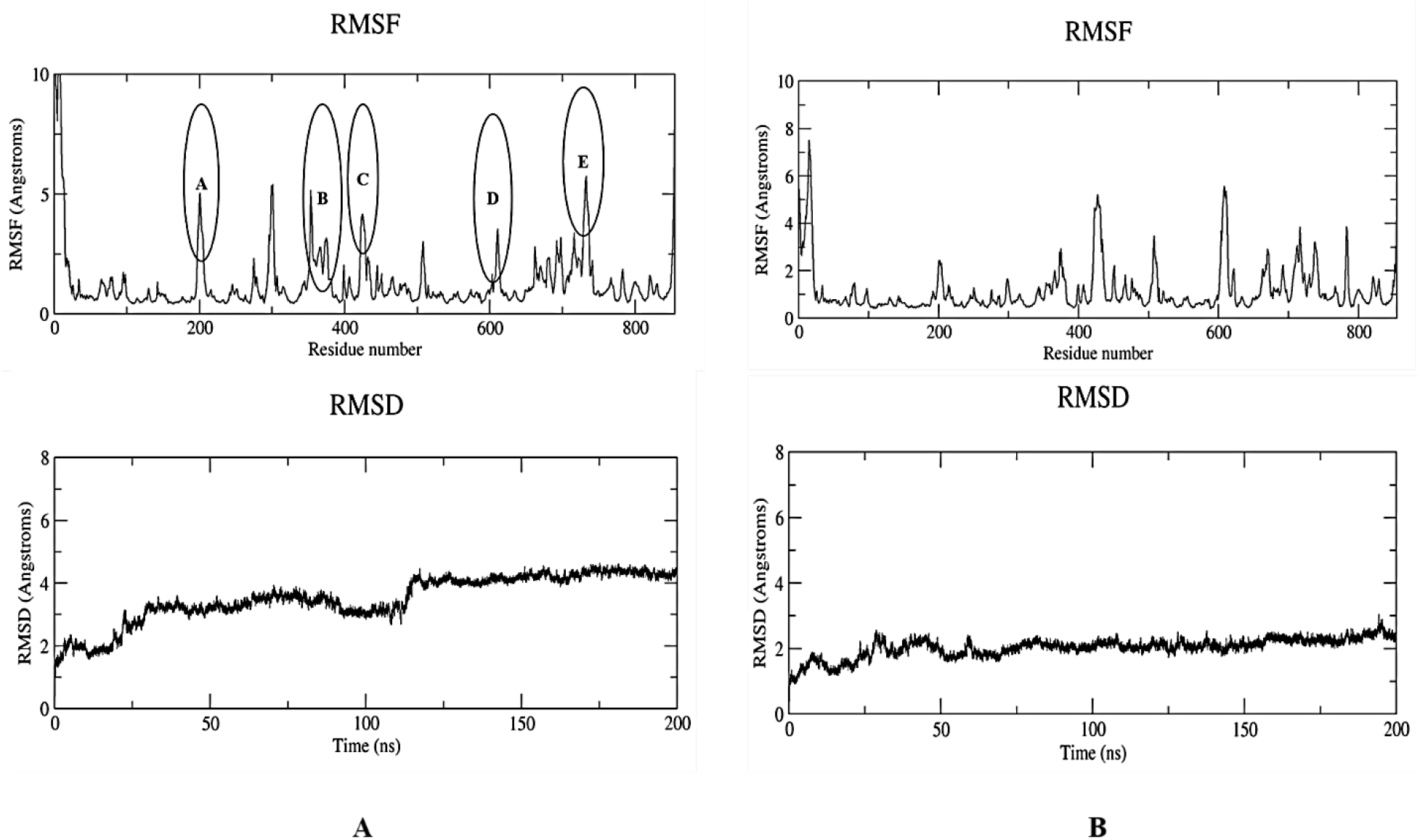
Molecular Dynamics (MD) simulation for *Pf*Bgl3A. Root mean square fluctuation (RMSF) and root mean square deviation (RMSD) values are shown before (A) and after (B) mutation. Regions marked in black circles are points with highest fluctuations selected for virtual mutation.

**Fig. 8.**
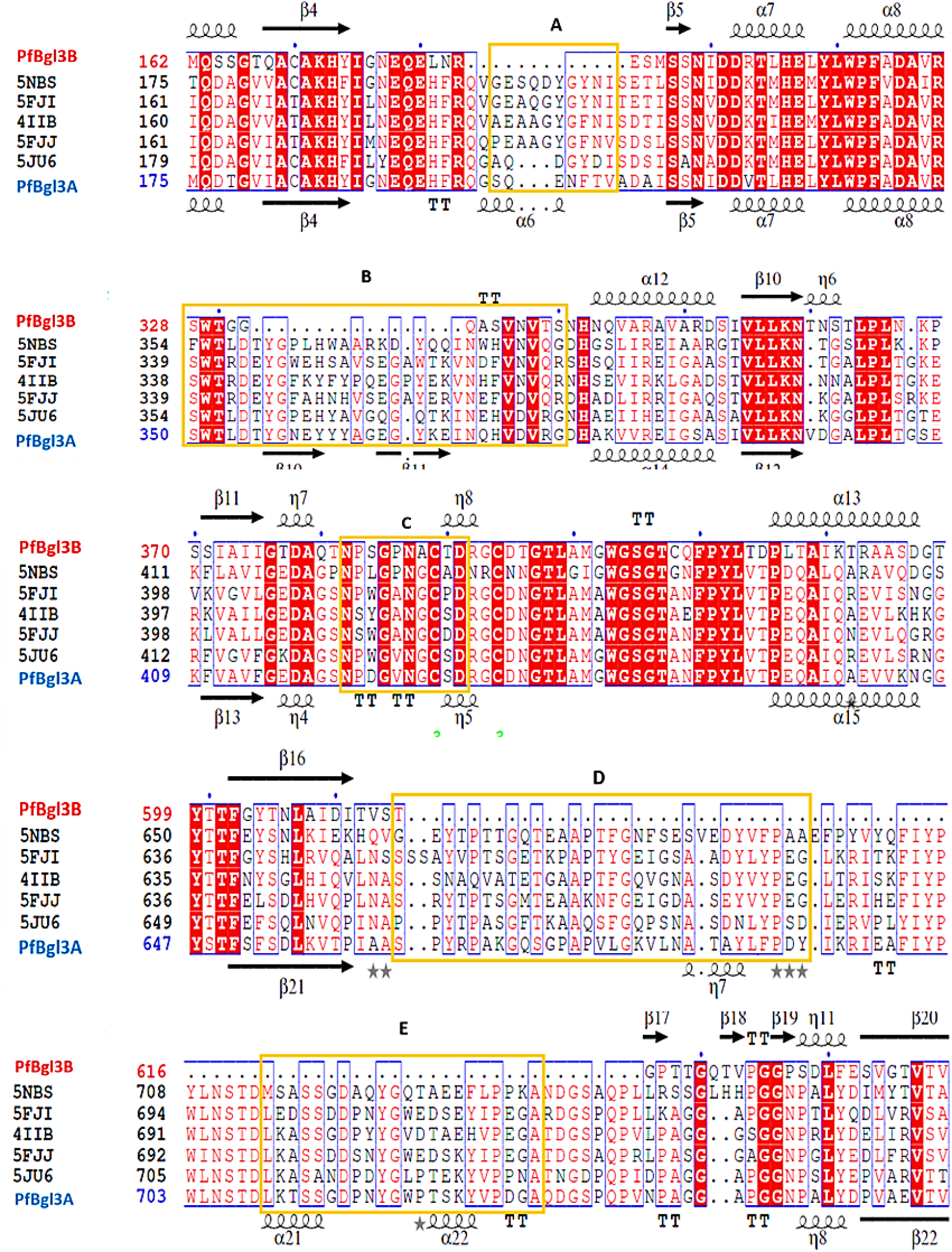
Multiple sequence alignment of extracellular GH3 β-glucosidases with GH3 β-glucosidases from *Neurospora crassa* (5NBS), *Aspergillus fumigatus* (5FJI), *Aspergillus aculeatus* (4IIB), *Aspergillus oryzae* and *Rasamsonia emersonii* (5JU6). Conserved regions are highlighted in red. Yellow boxes indicate regions corresponding to points with highest fluctuations (Fig. 6) where residues were selected for mutation.

**Fig. 9.**
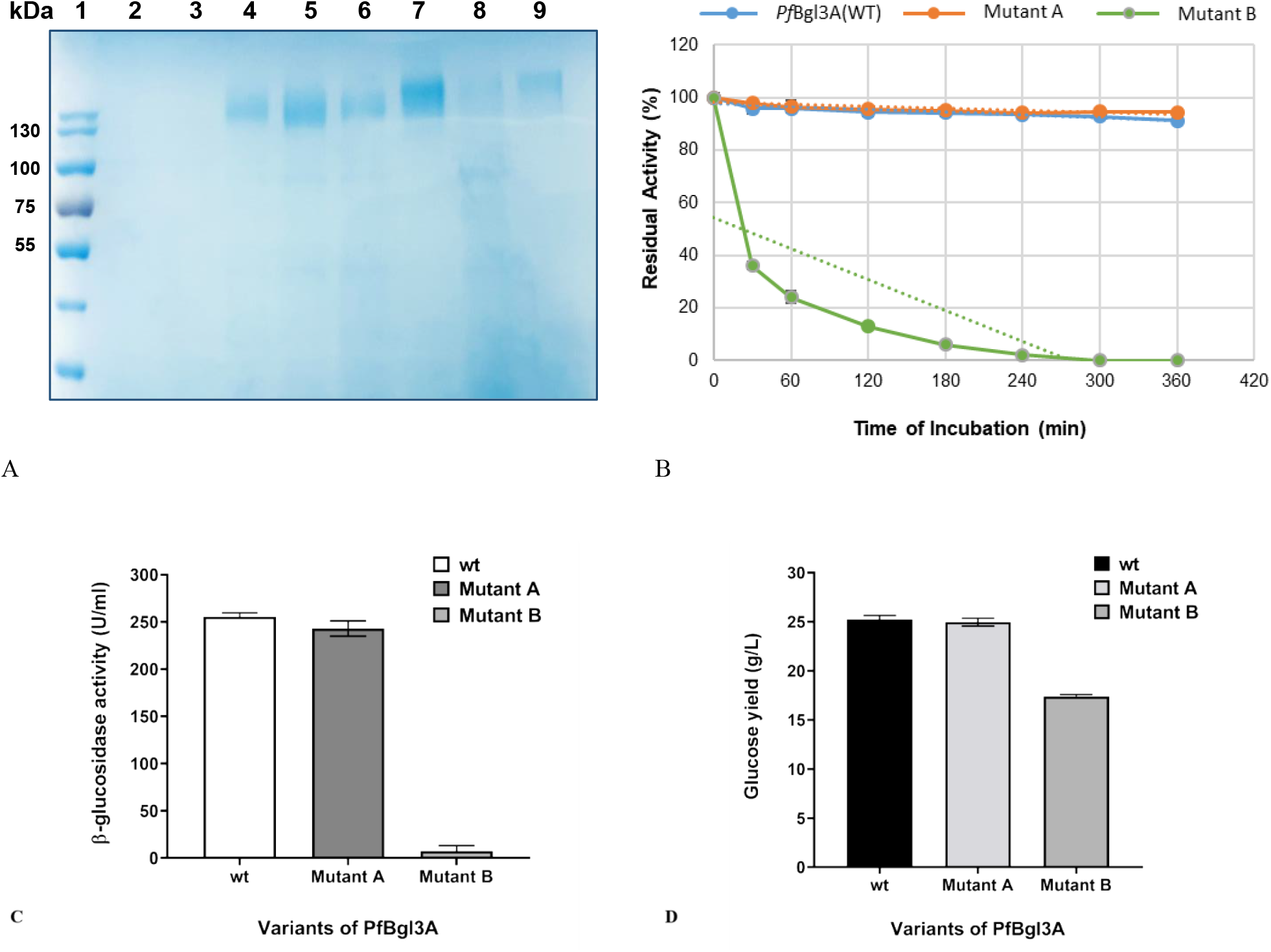
Analysis of expression of *Pf*Bgl3A and mutants in Pichia pastoris. (A) SDS-PAGE showing protein expression. Lane 1; molecular weight markers, lanes 2-3; *P. pastoris* X-33, lanes 4-5; *Pf*Bgl3A, lanes 6-7; Mutant A, lanes 8-9; Mutant B. (B) Thermostability analysis of expressed proteins. (C) The β-glucosidase activity of expressed proteins. (D) Glucose yield from cellobiose hydrolysis.

**Table 3.**
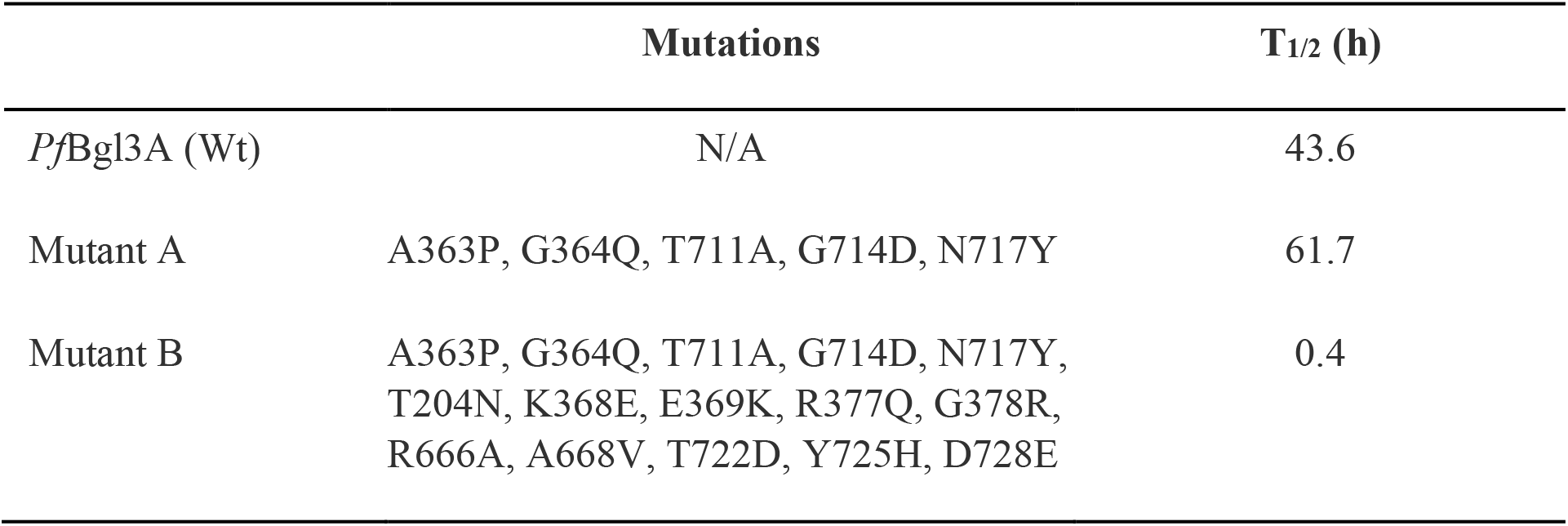
Half-life of *Pf*Bgl3A and its mutants after prolonged incubation at 50°C.

In conclusion, successful genetic and protein engineering strategies of *P. funiculosum* for improved biomass saccharification, requires an in-depth knowledge of the components that make up the saccharification machinery of the fungus. In this study, the β-glucosidase component was examined. The presence of multiple β-glucosidases was confirmed and their individual contribution to the overall hydrolyzing capacity of the secretome assessed. Additionally, a mutation strategy geared towards improved thermostability was performed. The results of this study gives an insight to the β-glucosidase profile of this hypercellulolytic fungus, highlights the significant contributory β-glucosidase and sets the foundation for future genetic interventions.

## MATERIALS AND METHODS

### Strains and culture conditions

The fungal strain used in this study was *Penicillium funiculosum* Mig1^88^ (PfMig1^88^) which is *Penicillium funiculosum* NCIM1228 having Migl catabolite repressor deleted (18). The culture media for sporulation of the fungus was LMP (1%Malt extract, 0.05% peptone and 1.5% agar). For transcript levels, LC-MS/MS analysis, and enzyme activity determinations, PfMig1^88^ was grown in modified Mandel’s media with Avicel and wheat bran (2%) as carbon source and in an optimized cellulase-inducing media (CIM) consisting of soya peptone (24 g/L), wheat bran (21.4 g/L), microcrystalline cellulose (24 g/L), KH_2_PO_4_ (12.4 g/L), K_2_HPO_4_ (2.68 g/L), (NH4)2SO4 (0.28 g/L), CaCO3 (2.5 g/L), corn steep liquor (1%), urea (0.52 g/L), and yeast extract (0.05 g/L). Final pH of the media was adjusted to 5.0 (38). The primary culture was prepared by inoculating potato dextrose broth (PDB) with 10^7^ conidiospores/ml from fungal cultures on LMP plates. This was incubated at 28 °C and 150 rpm in an incubator shaker (Innova 44, Eppendorf AG, Germany) for 36 h before being used to inoculate secondary media (50 ml of culture media in 250 ml Erlenmeyer flasks).

*Escherichia coli* DH5α was used for plasmid propagation and cloning while pCambia 1302 was used for construction of deletion cassettes. *Agrobacterium tumefaciens* LBA4404 strain was used for Agrobacterium-mediated transformation (AMT) of PfMig1^88^.

### Sequence and phylogenetic analysis of *Pf*Bgls

Genome sequencing and annotation of *Penicillium funiculosum* NCIM1228 had previously been done in our laboratory (16). The β-glucosidases identified were assigned names according to their respective families and their sequences deposited in GenBank (see supplementary material). Signal peptide prediction as well as conserved domain analysis was done with SignalP 6.0 Server (https://services.healthtech.dtu.dk/service.php?SignalP) and NCBI Conserved Domain Search (https://www.ncbi.nlm.nih.gov/Structure/cdd/wrpsb.cgi) respectively. NCBI BlastP (http://www.ncbi.nlm.nih.gov/) was used to find orthologues of *Pf*Bgl3A, *Pf*Bgl3B and *Pf*Bgl1A. Sequences were downloaded in Fasta format and multiple sequence alignment performed using Clustalx 2.1 software. The phylogenetic tree was constructed with MEGA 11 software (47) using neighbor-joining algorithms with 500 bootstrap replications.

### Homology protein structure modelling and docking

In order to find a few best templates for the homology modelling of BGLs, we ran HMMSEARCH (48) with protein sequences of β-glucosidases against the online available PDB database. The respective Bgls, and the protein sequences of the best suitable templates were aligned using a multiple sequence alignment tool, Clustal Omega. Based on the sequence similarities and alignment of the PDB templates with the Bgls, multiple PDB templates were chosen. (PDB IDs-“5FJI, 4IIB, 5FJJ and 5JU6” for *Pf*Bgl3A and *Pf*Bgl3B; “4JHO, 7F3A, 6M55, 3AI0 and 1V02” for *Pf*Bgl1A). These templates were used to perform homology modelling for each of the *Pf*Bgls using MODELLER version 9.23. Ten protein models were generated. The top-ranked models were selected (based on their Discrete Optimized Protein Energy model values) after performing quality assessments, including Ramachandran plots using PROCHECK.

For docking, each *Pf*Bgl protein and cellobiose were docked separately using Maestro tool of Schrodinger suite (49). The cluster representative protein structure for each β-glucosidases was used for the docking protocol. Firstly, protein structure was prepared using Protein Preparation Wizard of Schrodinger Suite, then hydrogen atoms were added, thereafter optimization of hydrogen bonds was performed and the protonated states of His, Gln, and Asn verified. An energy minimization step was subsequently done using OPLS4 force field. The binding site of the ligand in each selected model was taken as a reference for grid generation using Glide grid. Ligand preparation was done using LigPrep of Schrodinger Suite. In the final step, the ligand was docked to the proteins using “Extra precision mode” of Glide. Top-ranked conformers were selected based on their Glide score.

### Molecular dynamic simulations

We performed point mutations in *Pf*Bgl3A modelled protein structure using Rotamer program of UCSF Chimera (50) with Dunbrack 2010 library (51). The modelled protein structures of *Pf*Bgl3A and mutated variants (mutant A and mutant B) were used to perform molecular dynamic simulations separately using AMBER18 (52). Data analysis was done using AmberTools18 (53). Topology and parameter files for each protein was generated using leaprc.ff99SBxildn force field. The systems were prepared separately, first solvated using TIP3P water model with a buffer size of 12 Å in Octahedral box with Na^+^ and Cl^-^ions added to neutralize counter ions accordingly in their respective systems. The systems were equilibrated in multiple stages: firstly, protein was fixed and then water and ions were minimized, followed by the minimization of the entire system. The energy minimization procedure included initial 500 cycles of steep descent (SD) algorithm and then the remaining 19,500 cycles of conjugate gradient algorithm (CG) totaling 20,000 steps. In the next stage each system was heated-up from 0 to 303.15K over 100 picoseconds with a collision frequency of 2.0 picosecond ^-1^. Langevin thermostat (NTT=3) with weak restraints of 2 kcal mol^-1^ Å^-2^ on solute was used for temperature regulation. Next we performed density equilibration for 50 picoseconds and constant pressure equilibration at 5 nanoseconds. Finally, molecular dynamics trajectories were produced for 300 nanoseconds at 303.15K using PMEMD module of the AMBER18 suite. We used 2fs time step for all molecular dynamics simulation stages and all atoms involving hydrogen atoms were constrained using the SHAKE algorithm. For all the simulations, coordinates were saved in a single trajectory file every 10 picoseconds. All further analyses (RMSD, RMSF, and cluster analysis) were performed on the last 200ns molecular dynamic simulation trajectory by removing the initial first 100ns trajectory using the CPPTRAJ module (54) of AmberTools18.

### DNA/RNA isolation, cDNA synthesis and qRT-PCR

Mycelia of *Pf*Mig1^88^ was grown in PDB and harvested by filtration using Mira cloth. DNA was then isolated using the Quick-DNA Fungal/Bacterial Kit (Zymo Research USA).

For RNA isolation, mycelia was cultivated in modified Mandel’s media containing 2 g/l KH_2_PO_4_, 0.3 g/l CaCl_2_.2H_2_O, 0.3 g/l MgSO_4_.7H_2_O, 1.4 g/l (NH_4_)_2_SO_4_, 0.25 g/l Peptone, 0.1 g/l Yeast extract, 5 mg/l FeSO_4_, 1.6 mg/l MnSO_4_, 1.4 mg/l ZnSO_4_, 2 mg/l CoCl_2_.6H_2_O, 0.1 % Tween 80 and 4% carbon source. Mycelia was harvested and immediately flash frozen with liquid nitrogen. Total RNA was isolated with the Qiagen Plant mini RNeasy kit. Isolated RNA was treated with DNAse prior to cDNA synthesis which was carried out using the SuperScript™ III First-Strand Synthesis kit (Invitrogen). cDNA was stored at −80°C until used.

qRT-PCR was done using the iTaq Universal SYBR Green Supermix and CFX96 qPCR detection system (Bio-Rad). The reaction was performed in a 10 μl reaction volume containing 1 μl of cDNA, 5 μl of 2x SYBR Green, 2 μl of the primer pair and 2 μl of ddH_2_O. Primers (Table 1) were designed flanking two exons to avoid amplification of any genomic DNA contaminants. qRT-PCR was done in biological and technical replicates with actin and tubulin as endogenous controls. Relative expression levels were normalized to tubulin.

### LC-MS/MS analysis

Sample preparation for proteomic analysis was done as described in detail by Ogunyewo *et al*., (38). Secretome containing 50μg of protein was reduced and alkylated by dithiothreitol (DTT) and iodoacetamide (IAA) treatment and then digested with trypsin (16). The digested samples were desalted using C18 spin columns (Thermo Fisher Scientific) before being subjected to LC-MS/MS which was performed using Orbitrap Fusion Lumos Tribrid Mass Spectrophotometer (Thermo Fisher Scientific, Singapore).

### Deletion of *Pf*Bgl3A, *Pf*Bgl3B and *Pf*Bgl1A

Construction of deletion cassettes: The open reading frames of *Pf*Bgl3A, *Pf*Bgl3B and *Pf*Bgl1A with 500 bp upstream and downstream were amplified from the genome of PfMig1^88^ through PCR using forward and reverse *Pf*Bgl primer sets. (Table 1). The amplicons were first cloned into a pJET blunt vector containing an ampicillin resistance gene. They were then removed from the shuttle vector by restriction digestion and cloned into pCambia1302 at the XhoI/MauBI restriction sites between the left and right T-DNA borders. The β-glucosidase deletion cassettes for *Pf*Bgl3A, *Pf*Bgl3B and *Pf*Bgl1A were constructed by replacing 2090bp, 1779bp and 2178bp region respectively at the middle of the ORF with a 1.9 kb hygromycin gene by restriction digestion and ligation. The hygromycin insert previously obtained in the laboratory (21) was amplified from pARI vector through PCR using Hyg primers (Table 1).

Bacterial cloning and transformation: The deletion vectors were transformed into *E. coli* DH5α. Transformants were selected with 50 μg/ml kanamycin. Plasmids isolated from positive transformants were used to transform *Agrobacterium tumefaciens* LBA4404 through the heat shock method as described by Randhawa et al. (21).

Agrobacterium-mediated transformation of *P. funiculosum:* Positive transformants were selected with 50 μg/ml kanamycin and 30 μg/ml rifampicin. These were then pre-induced for T-DNA mobilization with induction media (55) until OD_600_ 0.3 was obtained. Equal volumes of the preinduced cells and freshly harvested PfMig1^88^ spores were incubated at 22°C for 1 h and then plated on cellophane membrane covered IM agar plates containing acetosyringone. The plates were incubated for 48 h until a whitish mycelial mass appeared on the cellophane membranes. The membranes were then transferred to selection plates containing 500mM cefotaxime, 0.1 mM chlorpromazine, 0.007 % Triton-X and 100 μg/ml hygromycin. The plates were subsequently incubated at 28°C until transformants appeared.

### Design and expression of thermostable variants of *pf*Bgl3A in *Pichia pastoris*

From homology modelling data, root mean square deviation (RMSD) and root mean square fluctuation (RMSF) values were calculated and points of highest fluctuations were noted. Multiple sequence analysis was then performed with sequences of thermostable β-glucosidases which had structures in PDB. From the data obtained, two mutants A and B were designed. Mutant A had amino acid changes with > 2-fold change in RMSD values while mutant B had all mutations in A and additional mutations with ≤ 2-fold change in RMSD (Supplementary Table S3).

Nucleotide sequences of *Pf*Bgl3A, mutant A and mutant B without signal peptides were synthesized by GenScript USA, Inc. (New Jersey, USA) and cloned into the pPICZαA vector at the *XhoI/XbaI* sites. The vectors, designated as pPICZαAPfBgl3A, pPICZαAPfBgl3A_A and pPICZαAPfBgl3A_B, respectively, were transformed into *E. coli* DH5α competent cells for multiplication, after which plasmids were isolated using Qiagen mini-prep kit and linearized with *SacI* restriction endonuclease enzyme. The linearized plasmids were subsequently transformed into competent *Picha pastoris* X-33 cells by the method described by Kumar et al., (56). After transformation, the mixture was plated on YPD agar plates containing 100 μg/ml zeocin, for the selection of positive transformants. The transformants were further screened by reselection on YPD agar plates containing 1000 μg/ml zeocin for 48 h and polymerase chain reaction (PCR) with the AOXI521F and AOX1-R primer pairs (Table 1). The selected positive transformants were then inoculated into buffered glycerol-complex medium (BMGY) for 24 h and transferred into methanol-complex medium (BMMY) for β-glucosidase expression with the addition of 1% methanol every 12 h until the end of the fermentation process. Induction of the expressed protein was monitored by collecting culture samples every 24 h and analyzed by sodium dodecyl sulfate-polyacrylamide gel electrophoresis (SDS-PAGE).

### Analytical assays

MUG-zymogram analysis was used for in-gel detection of β-glucosidase activity as described by Li et al., (37) with some modifications. Samples were analyzed by native PAGE with 10% and 5% polyacrylamide separation and stacking gels, respectively.

The β-glucosidase activity was determined in 50mM sodium citrate buffer pH 4.0 at 50 °C using 4-nitrophenyl β-D-glucopyranoside (pNGP) as substrate (18, 19). The amount of p-nitrophenol released from pNPG was measured and one unit of activity was defined as the amount of enzyme required to release 1 μmol pNP min^-1^ under standard assay conditions. Protein in samples was estimated by the bicinchoninic acid (BCA) method using bovine serum albumin (BSA) as standard.

### Biomass Hydrolysis

Saccharification efficiency on sodium hydroxide treated sugarcane bagasse (SCB) was determined as previously described by Ogunyewo et al., (19). Saccharification was performed in 50 ml falcon tubes in an incubator shaker at 50 °C for 96 h. the reaction mixture contained sodium hydroxide-treated SCB at 15 % biomass load and 5FPU/g biomass enzyme load.

### Thermostability Assessment

Thermostability was determined by incubating the enzyme at 50 °C for 24-120 h and then the β-glucosidase activity was measured using pNPG as substrate. Results are presented as a residual activity where the activity of the enzyme at T_0_ is defined as 100% (45).

### Cellobiose hydrolysis

The cellobiose hydrolyzing efficiency of the mutants was evaluated with the method described by Zhang et al. (57). The reaction was set up with 15 mM cellobiose in 50 mM citrate phosphate buffer pH 4.8 as substrate. One microliter of appropriately diluted enzyme solution was temperate to 50°C. One microliter of the substrate was then added. The reaction mixture was then incubated for exactly 30 min. The reaction was stopped by boiling at 100 °C for 5 min. The amount of glucose released was measured by high-performance liquid chromatography (HPLC) with an Aminex HPX-87H anion-exchange column (Bio-Rad, USA) and a refractive index (RI) detector to analyze the released glucose anion-exchange chromatography. The filtered mobile phase (4 mM H_2_SO_4_) was used at a constant rate of 0.3 ml/min with the column, and the RI detector temperature was maintained at 35 °C. The concentration of glucose was calculated from the calibration curve of the glucose standard.

### Data and statistical analysis

All experiments were performed in triplicates and results are presented as means and standard deviations. Data was collected on a Microsoft Excel spreadsheet on which the average and standard error of means were calculated. Graphs were created with Graphpad Prism 8.0 software. One-way analysis of variance (ANOVA) was used to test statistical significance where appropriate.

## AUTHOR CONTRIBUTIONS

SSY and OEO conceived and designed the study. MG performed the computational analysis, OEO, OAO and KS conducted the experiments. OEO and MG wrote the manuscript. SSY supervised the entire study. All authors read and approved the final manuscript.

## ACKNOWLEDGEMENTS

This study was funded by DBT, Government of India via Grant no. BT/PR/Centre/03/2011-Phase II. OEO is thankful to International Centre for Genetic Engineering and Biotechnology (ICGEB) for funding through the Arturo Falaschi postdoctoral fellowship. The authors thank Mr Girish H. Rajacharya for HPLC sample analysis.

